# Assembly-pathway regulation dictates pH-responsive actuation in the R-body protein machinery

**DOI:** 10.64898/2026.05.21.725848

**Authors:** Koki Date, Kosuke Kikuchi, Tatsuya Niwa, Hironari Kamikubo, Sota Masumura, Thuc Toan Pham, Keiichi Okisawa, Hideki Taguchi, Takafumi Ueno

## Abstract

Refractile bodies (R-bodies) are protein assemblies that form tightly rolled morphologies in cells and undergo large-scale extension into long spirals in response to environmental stimuli. The type 51 R-body, the focus of this study, is assembled from four Reb proteins and undergoes a rapid repeatable ∼50-fold extension. Although R-bodies were first described more than 70 years ago, it has remained unclear how RebC and RebD contribute to the formation and function of this four-protein machinery even though RebA and RebB have been proposed as its major components. We characterized the wild-type R-body (Rb_WT) and a series of *reb* gene knockout mutants by combining in-cell, biochemical analyses, small-angle X-ray scattering (SAXS) and attenuated total reflection Fourier-transform infrared (ATR-FTIR) spectroscopy. We found that RebD is incorporated into Rb_WT as a minor component, while RebC is not detectably incorporated into the final assembly. Mutants lacking either RebA or RebB still form roll-like assemblies but lack pH-dependent extension. In contrast, mutants lacking RebC and/or RebD exhibit a substantially lower propensity for roll formation and instead undergo off-pathway aggregation with increased β-sheet content. SAXS analyses further indicate that roll-like morphology does not ensure formation of the ordered lamellar architecture characteristic of functional Rb_WT. Thus, pH-responsive R-body actuation is dictated not simply by the major proteins that constitute the final architecture, but by a regulated assembly pathway that builds the ordered lamellar architecture required for actuation. These findings establish assembly-pathway regulation as a key principle for constructing dynamic protein architectures capable of stimuli-responsive mechanical actuation.

## Introduction

Protein assemblies achieve ordered, functional mesoscale architectures through the coordinated contributions of proteins that play distinct molecular roles (1). In systems such as viral capsids, clathrin coats, bacterial curli, and microtubules, some components mainly constitute the mature architecture, whereas others guide nucleation, stabilize assembly intermediates, or provide templates for higher-order packing to direct productive assembly across length scales (2–6). Although recent advances in computational protein design have enabled the creation of programmable heteromeric assemblies (7–10), current strategies focus on stable target architectures rather than on molecular roles that guide assembly pathways. In dynamic, stimuli-responsive assemblies, large-scale morphological transformation depends on the ordered packing and metastable structural states established through productive assembly pathways (11, 12). Consequently, limited understanding and control of assembly pathways constrain the engineering of protein assemblies capable of such transformations (13, 14).

Refractile bodies (R-bodies) are microscale protein assemblies that undergo stimuli-responsive, pronounced morphological transformations (Fig. 1*A*-*C*, ref. 15). Expressed in endosymbionts of *Paramecium*, they adopt a tightly rolled ribbon morphology with a ∼0.4 μm diameter at neutral pH (16, 17). When the environmental pH decreases, the R-bodies extend into a ∼20 μm spiral ribbon within one second (Fig. 1*A-C*, Movies S1 and S2, ref. 16, 17). This rapid extension triggers cell death in susceptible strains, even though R-bodies themselves are non-toxic (18–22). Previous studies demonstrated that type 51 R-body can be heterologously expressed and assembled in *Escherichia coli* (Fig. 1*D*, ref. 23), broadening further fundamental investigations and engineering applications. Despite the long-standing interest in R-bodies, a systematic structural characterization of their formation pathways and the specific roles of individual Reb proteins has been lacking (24– 26).

**Figure 1.**
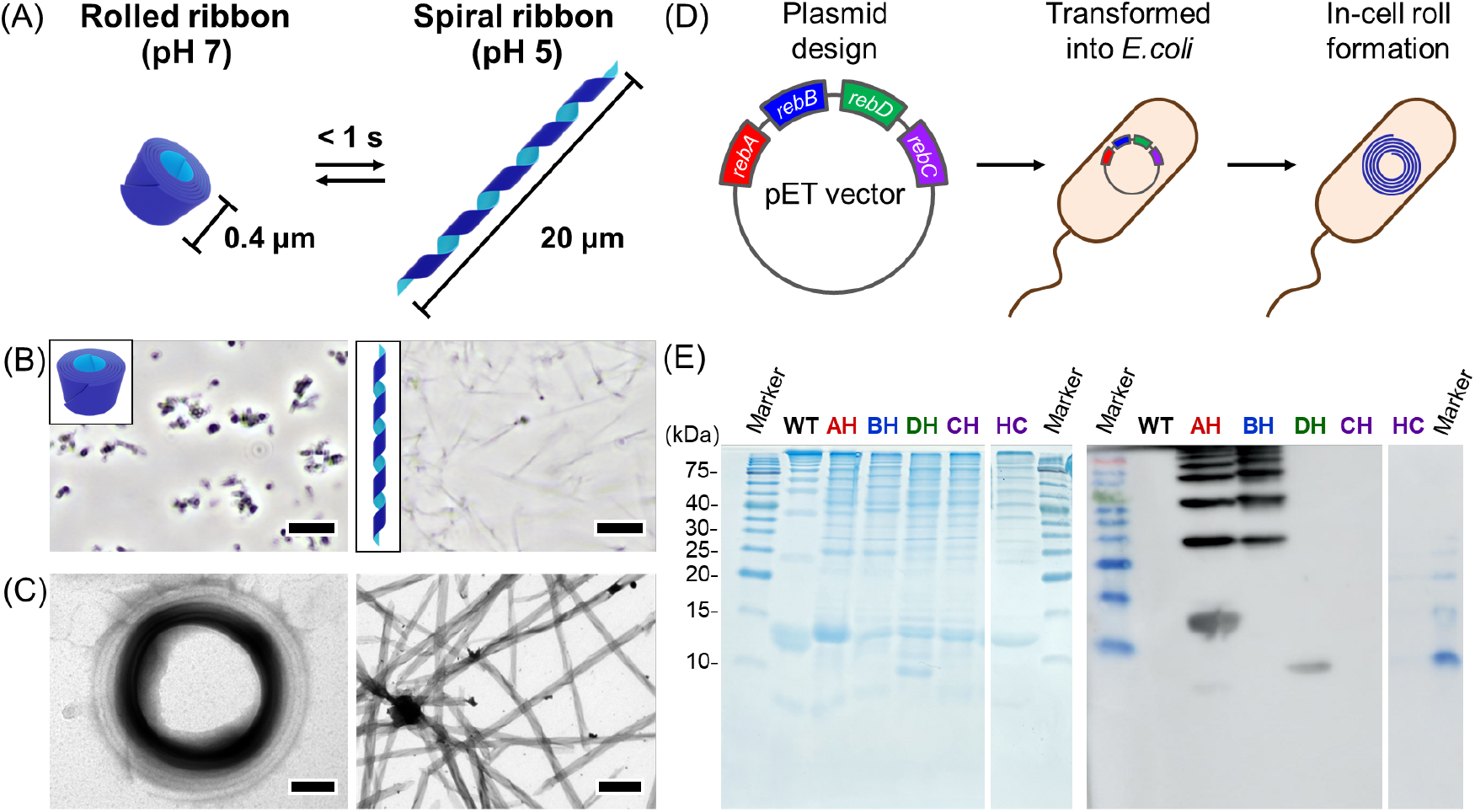
pH-dependent transformation and composition of Rb_WT. (A) Schematic illustration of pH-dependent reversible transformation between the roll (contracted) and spiral (extended) morphologies. (B) Phase-contrast microscopy images of isolated Rb_WT at pH 7.0 (contracted, left) and after buffer exchange to pH 5.0 (extended, right). (C) TEM images of isolated Rb_WT at pH 7.0 (contracted, left) and after buffer exchange to pH 3.0 (extended, right). (D) Schematic illustration of in-cell roll formation. (E) SDS-PAGE and the corresponding Western blotting analysis of isolated His-tagged mutants. AH, BH, DH, and CH represent **Rb_WT(RebX-His)**, where X = A, B, D, or C. HC represents **Rb_WT(His-RebC**). Scale bars: 5 μm (B), 100 nm (C, left), 1 μm (C, right).

The formation of type 51 R-body is governed by a four-gene operon comprising *rebA, rebB, rebC*, and *rebD* (Fig. 1*D*, ref. 15, 27, 28). Each gene encodes its respective Reb proteins (RebA, RebB, RebC, and RebD). RebA and RebB have been proposed as the major components (23, 27), but this assignment is supported only by electrophoretic experiments, in the absence of any structure-based characterization. Subsequent studies on R-bodies have built upon this limited framework, and comprehensive identification of protein contributions at the molecular and structural levels has remained unexplored. Thus, elucidating how each Reb protein contributes to establishing the extensible ribbon architecture stands as a central question in the R-body assembly process.

To clarify the contributions of individual Reb proteins, we expressed and characterized the wild-type R-body (Rb_WT) and a series of single-, double-, and triple-gene knockout mutants. We analyzed their protein compositions, in-cell morphologies, isolated morphologies, pH-responsiveness, nanoscale structural features, and secondary structures. Although rolled morphologies were observed in some mutants, the pH-responsive extension was only observed in Rb_WT, indicating that all four Reb proteins are essential for functional R-body assembly. Our results reveal that RebA and RebB are major structural components, RebD is a minor structural component which is essential for roll formation, and RebC promotes roll formation as an assembly factor, favoring an α-helix-rich lamellar-ordering pathway over β-sheet-rich off-pathway aggregation. Together, these findings reveal that pH-responsive R-body actuation is not determined by roll-like morphology or major structural components alone, but by a regulated assembly pathway that establishes the ordered lamellar architecture required for the complex functionality of this system.

## Results

### SDS-PAGE analysis of Rb_WT

The expression and isolation of Rb_WT were performed based on a previous report (see *SI Appendix*, Methods, ref. 17). The SDS-PAGE analysis of Rb_WT showed characteristic ladder bands ranging from ∼10 kDa to over 245 kDa, as reported previously (Fig. 1*E*, ref. 23, 27). To assign these bands, we performed Western blotting using His-tagged mutants (**Rb_WT(RebX-His)**, where X = A, B, C, and D), in which an oligo-histidine tag was fused to the C-terminus of each Reb protein (Fig. 1*E*, and *SI Appendix*, Table S1). **Rb_WT(RebA-His)** and **Rb_WT(RebB-His)** showed ladder bands beginning at ∼13 kDa and ∼23 kDa, respectively. **Rb_WT(RebD-His)** showed only a single band below 10 kDa. The corresponding band in the CBB-stained gel comprised less than 3% of the total lane intensity in Rb_WT, suggesting that RebD is a minor component of Rb_WT (*SI Appendix*, Fig. S1). RebC-His was not detected by either SDS– PAGE or Western blotting when using **Rb_WT(RebC-His)**. Furthermore, **Rb_WT(His-RebC)**, in which an oligo-histidine tag was fused to the N-terminus of RebC, did not show the bands of His-RebC by SDS–PAGE and Western blotting analyses, while singly expressed His-RebC was detected by both analyses (*SI Appendix*, Fig. S2). These results suggest that RebC is not incorporated into Rb_WT in a detectable amount.

### Construction of single, double-, and triple-gene knockout mutants

To clarify the contributions of each Reb protein in the R-body formation, we constructed a series of *reb* gene knockout mutants, comprising single-, double-, and triple-gene knockouts, denoted by the retained Reb components (e.g., **Rb_ABD** indicates a RebC-knockout mutant). The knockout mutants were expressed using the same procedure as for Rb_WT (see *SI Appendix*, Materials and Methods).

The intracellular assemblies of the knockout mutants were observed by TEM on chemically fixed ultrathin sections (Fig. 2*A*). When Rb_WT was expressed, 60% of cells harbored one or two roll assemblies of Rb_WT. Mutants retaining both RebD and RebC, namely **Rb_BDC** (49%), **Rb_ADC** (42%), and **Rb_DC** (67%), similarly showed frequent roll formation (Fig. 2*B*). In contrast, mutants lacking RebC and/or RebD exhibited a substantially lower propensity for the roll formation. Notably, the roll formation was found to be markedly more frequent in **Rb_ADC** and **Rb_BDC** than in **Rb_AD** (6%) and **Rb_BD** (1%), respectively (Fig. 2*B*), indicating that the presence of RebC strongly promotes the roll formation, although RebC itself was not detected in the R-body (Fig. 1*E*). Thus, RebD and RebC play a central role in enabling intracellular roll formation, whereas their absence leads to markedly reduced frequency of detectable roll assemblies within cells.

**Figure 2.**
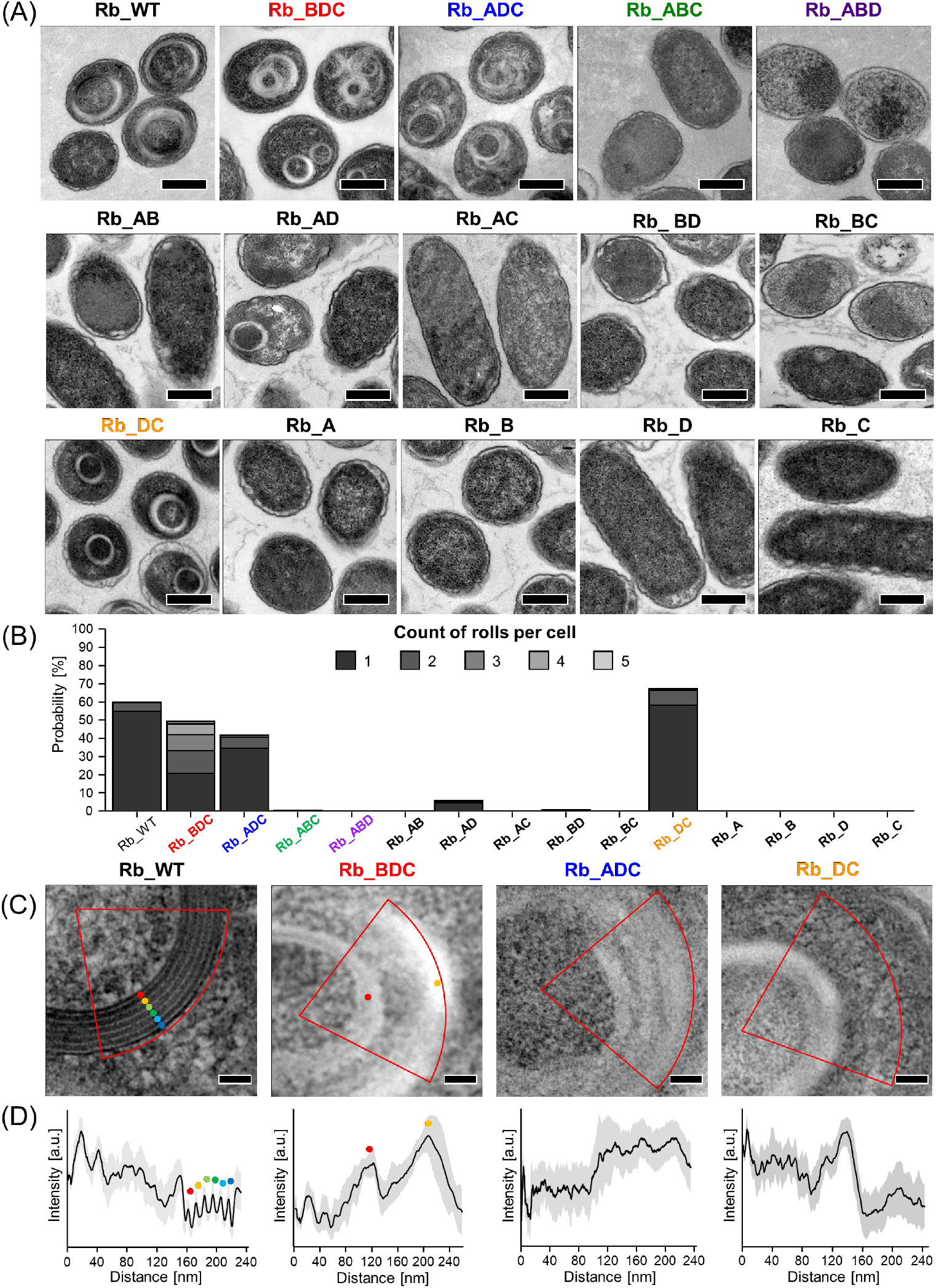
In-cell TEM analysis of Rb_WT and knockout mutants. (A) Representative TEM images of chemically fixed ultrathin sections. (B) Quantification of roll formation frequency in each mutant. (C) Magnified view of the selected strains exhibiting roll assemblies. (D) Circularly averaged radial contrast profiles (black lines) with corresponding standard deviations (shaded areas). Colored circles indicate the correspondence between panels (C) and (D). Scale bars, 400 nm (A), 50 nm (C).

Rb_WT roll assemblies exhibit clear lamellar periodicity in layer-to-layer contrast, whereas **Rb_BDC, Rb_ADC**, and **Rb_DC** did not show such well-defined patterns (Fig. 2*C*). Radial contrast analysis (see *SI Appendix*, Materials and Methods) revealed pronounced periodic peaks for Rb_WT, whereas no distinct peaks were observed for the mutants (Fig. 2*D*). The layer-to-layer periodic spacing in Rb_WT was determined to be 10.5 ± 0.3 nm based on the peak-to-peak distances in the radial contrast profiles (*SI Appendix*, Fig. S3). These results indicate that Rb_WT is characterized by a well-defined lamellar periodicity, which is not achieved solely by the formation of roll assemblies of the knockout mutants.

### Characterization of isolated knockout mutants

We isolated single-gene knockout mutants (**Rb_BDC, Rb_ADC, Rb_ABC**, and **Rb_ABD**), along with **Rb_DC**, which exhibited frequent intracellular roll formation, following the same procedure applied to Rb_WT (see *SI Appendix*, Materials and Methods). Notably, none of the knockout mutants showed detectable pH-dependent extension (*SI Appendix*, Fig. S4). The negative-stain TEM observation revealed that isolated Rb_WT, **Rb_BDC**, and **Rb_DC** retained rolled morphologies after isolation, whereas **Rb_ADC** appeared as fragmented sheet-like structures, and **Rb_ABC** and **Rb_ABD** exhibited globular aggregates (Fig. 3*A*, and *SI Appendix*, Fig. S5). These results indicate that the extensibility of the R-body is not an inherent aspect of the rolled morphology alone but emerges only when all four Reb proteins are present.

**Figure 3.**
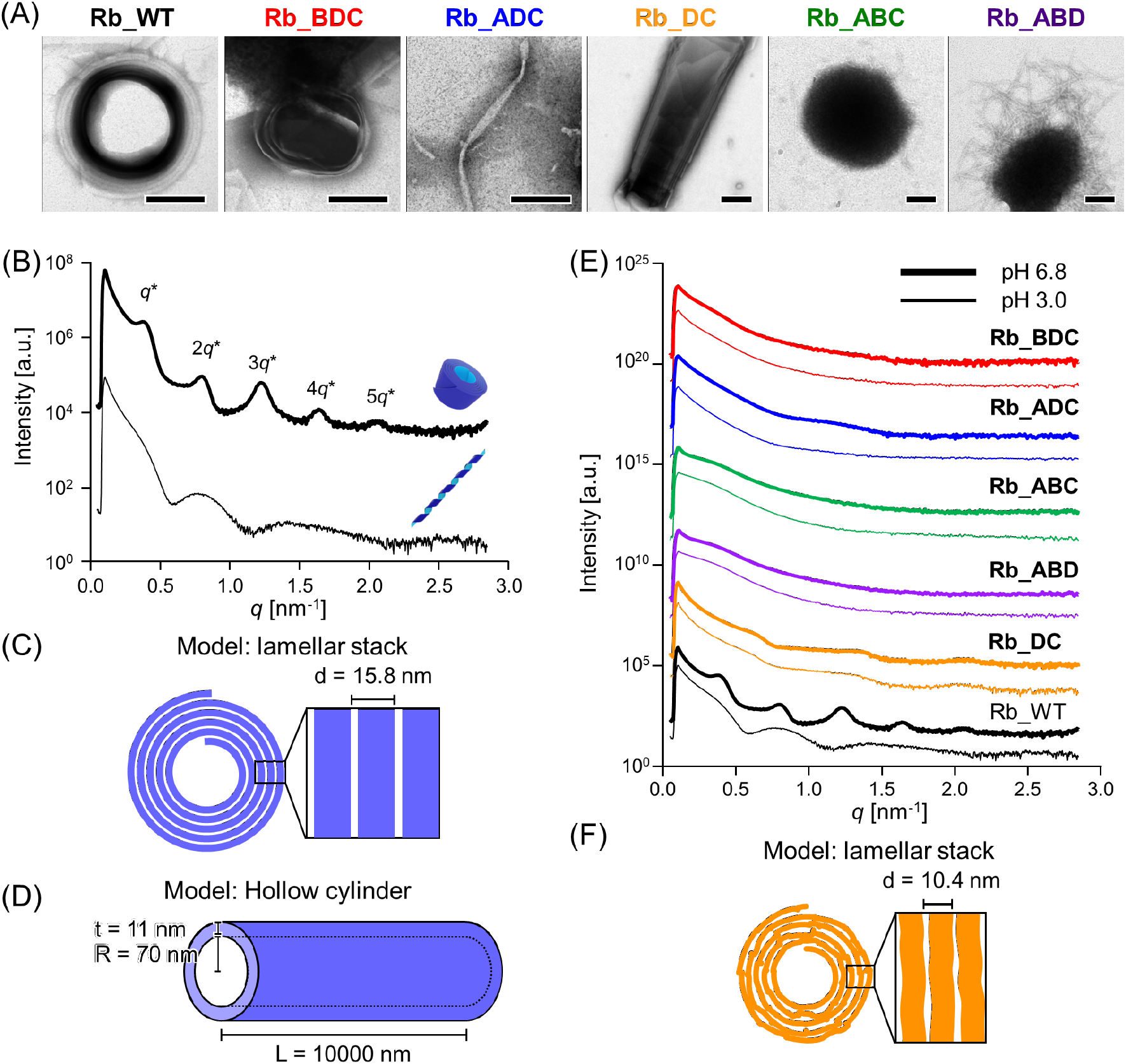
TEM observations and SAXS analyses of isolated assemblies. (A) Representative negative-stain TEM images of Rb_WT and knockout mutants at pH 7.0. Scale bars, 200 nm. (B) SAXS profiles of isolated Rb_WT at pH 6.8 (in a rolled morphology, thick lines) and pH 3.0 (in a spiral morphology, thin lines). The spiral state exhibits broad scattering features, whereas the rolled morphology is characterized by distinct Bragg diffraction peaks. (C) Schematic representations of the structural models used to simulate the theoretical lamellar periodicity in the rolled morphology of Rb_WT. (D) Schematic representations of the structural models used to simulate the theoretical sheet thickness in the spiral morphology of Rb_WT. (E) SAXS profiles of isolated Rb_WT and knockout mutants measured at pH 6.8 (citrate–phosphate buffer; thick lines) and pH 3.0 (citrate– phosphate buffer; thin lines). (F) Schematic representations of the structural models of **Rb_DC**.

SDS-PAGE analyses of **Rb_BDC, Rb_ADC, Rb_ABC, Rb_ABD**, and **Rb_DC** did not exhibit the characteristic ladder bands observed for Rb_WT (*SI Appendix*, Fig. S6). RebA and RebB were clearly detected as designed, whereas RebD was detected only in the presence of RebC, and the band corresponding to RebC (12.5 kDa) was not detected in any of the mutants. Instead, all mutants displayed multiple bands above approximately 8 kDa, suggesting the presence of additional protein components. LC-MS analysis revealed a high abundance of chaperone proteins, including IbpA, IbpB, ClpB, and HtpG, which are known to be overexpressed under protein aggregation stress in *E. coli* (*SI Appendix*, Fig. S7, ref. 29–32). Compared with its abundance in Rb_WT, IbpB was markedly enriched in **Rb_ABC** (∼99-fold) and **Rb_ABD** (∼110-fold), while more moderate increases were observed in **Rb_BDC** (∼13-fold) and **Rb_ADC** (∼15-fold) (*SI Appendix*, Fig. S7). IbpB was decreased but within the same magnitude observed in **Rb_DC** (∼0.32-fold) (*SI Appendix*, Fig. S7). These trends are consistent with the aggregation behavior observed by TEM (Fig. 3*A*). Together, these results suggest that, despite undergoing the same isolation procedure as Rb_WT, the single-gene knockout mutants that lack RebC or RebD contain substantial impurities and induce intracellular aggregation stress in *E. coli*.

### SAXS analyses of isolated mutants

To elucidate the structural features of this system, we performed small-angle X-ray scattering (SAXS) measurements (Fig. 3*B*). The characteristic peaks were observed in isolated Rb_WT in a rolled morphology at pH 6.8 (Fig. 3*B*). The SAXS profile of Rb_WT at pH 6.8 exhibited multiple Bragg peaks (Fig. 3*B*, bold line), which indicate a lamellar structure model with a periodic spacing of 15.8 nm (Fig. 3*C*, and *SI Appendix*, Fig. S8*A* and *B*). The discrepancy between this SAXS-derived interlamellar distance and the TEM-derived layer-to-layer distance (10.5 ± 0.3 nm, Fig. 2*D*, and *SI Appendix*, Fig. S3) is attributed to dehydration during TEM sample preparation (33). In a spiral morphology at pH 3.0, the SAXS profile of Rb_WT exhibited broad scattering features (Fig. 3*B*, thin lines), which match a hollow cylinder model with a sheet thickness of 11 nm (Fig. 3*D*, and *SI Appendix*, Fig. S8*C* and *D*). Thus, SAXS clearly distinguishes the roll and spiral morphologies of Rb_WT, revealing that only the rolled morphology exhibits diffraction peaks indicative of well-defined nanoscale lamellar order.

Conversely, no diffraction peaks were observed for the knockout mutants of **Rb_BDC, Rb_ADC, Rb_ABC**, and **Rb_ABD**, either under in-cell or isolated conditions (Fig. 3*E*, and *SI Appendix*, Fig. S9). The absence of SAXS diffraction peaks for **Rb_BDC** and **Rb_ADC**, despite their rolled morphologies observed by TEM (Fig. 2*A*, and Fig. 3*A*), demonstrates that the rolled morphology alone is insufficient to generate Bragg diffraction. **Rb_DC** showed a weak lamellar diffraction signature with a periodic spacing of 10.4 nm (Fig. 3*E* and *F*, and *SI Appendix*, Fig. S10*A* and *B*), smaller than that of Rb_WT (15.8 nm, *SI Appendix*, Fig. S8*A* and *B*). However, the quantitative analyses based on diffraction peak width revealed the cumulative lamellar disorder in **Rb_DC** (*SI Appendix*, Fig. S10*C*, ref. 34). Therefore, clear lamellar periodicity is observed only in Rb_WT, indicating that the co-existence of four kinds of Reb proteins is required for ordering the nanoscale structure.

### Secondary structure characterizations

To determine whether these nanoscale differences in lamellar organization originate from differences in secondary structure, we examined Rb_WT and the knockout mutants by attenuated total reflection Fourier-transform infrared (ATR-FTIR) spectroscopy and a Thioflavin T (ThT) assay. The spectra of Rb_WT at pH 7.5 and 5.0 exhibited strong absorption bands near 1650 cm^-1^ (Fig. 4*A*). These results suggest that α-helices are the predominant secondary structure across both pH conditions, consistent with a previous study using circular dichroism (CD) spectroscopy (17). Similarly, **Rb_DC** also showed high α-helical content (Fig. 4*B*). The ThT assay of Rb_WT and **Rb_DC** showed lower fluorescence intensity than other knockout mutants, indicating minimal β-sheet–rich amyloid-like assemblies among them (Fig. 4*C*). Since both **Rb_WT** and **Rb_DC** are characterized by rolled morphologies with lamellar-derived diffractions (Fig. 3*B* and *E*), these results suggest that the ordered rolled morphologies are formed dominantly by α-helical conformations.

**Figure 4.**
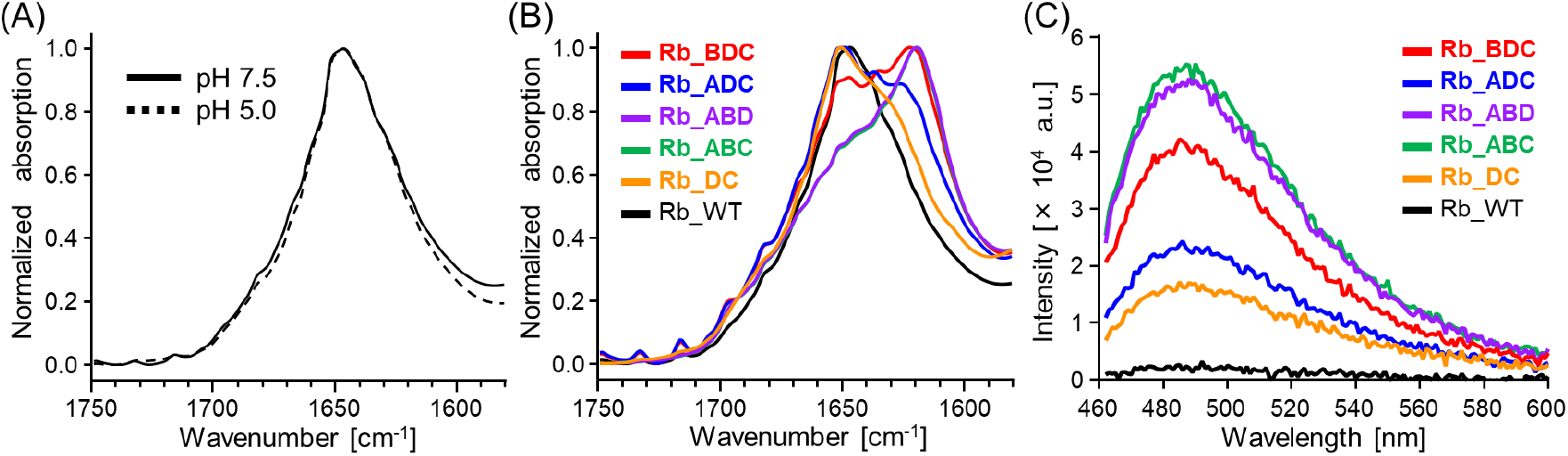
ATR-FTIR analysis of Rb_WT and knockout mutants. (A) The spectra of Rb_WT at pH 7.5 (solid line) and pH 5.0 (dashed line). (B) The spectra of isolated knockout mutants (**Rb_BDC, Rb_ADC, Rb_ABC, Rb_ABD, Rb_DC**) at pH 7.5. (C) Thioflavin T (ThT) assay of Rb_WT and knockout mutants. Emission spectra (λ_ex_ = 440 nm) of Rb_WT and knock-out mutants suspended in 50 mM Tris-HCl at pH 7.5. ThT fluorescence measurements were corrected by subtracting the background fluorescence of ThT measured in the absence of protein under identical buffer conditions. Knockout mutants exhibited increased ThT fluorescence intensity compared with Rb_WT, indicating amyloid-like β-sheet formation.

In contrast, the mutants lacking RebA or RebB (**Rb_BDC** and **Rb_ADC**) exhibited increased β-sheet content in ATR-FTIR measurements and enhanced fluorescence in the ThT assay (Fig. 4*B* and *C*). These analyses suggest that **Rb_BDC** roll assemblies and **Rb_ADC** fragmented sheets harbor amyloid-like β-sheet structures that are accessible to ThT binding. The mutants lacking RebD or RebC (**Rb_ABC** and **Rb_ABD**), which formed globular aggregates as observed in TEM (Fig. 3*A* and *SI Appendix*, Fig. S5), likewise showed higher β-sheet content and ThT fluorescence. These results suggest that RebD and RebC cooperatively inhibit amyloid-like β-sheet aggregation.

## Discussion

By constructing single-, double-, and triple-gene knockout mutants, this work clarified the distinct contributions of RebA–D in R-body formation and function. ATR-FTIR revealed that Rb_WT primarily consists of α-helical structures (Fig. 4*A*), which is rare for two-dimensional protein assemblies except for a few reports (35, 36). Western blotting analyses suggested RebA and RebB are the major structural components of Rb_WT, whereas RebD is a minor yet essential structural component for roll formation (Fig. 1*E*, Fig. 2*A* and *B*, and *SI Appendix*, Fig. S1). Although RebC was not detected as a structural component in the isolated R-body, its expression was essential for efficient intracellular roll formation (Fig. 1*E*, and Fig. 2*A* and *B*). The combination of RebD and RebC (**Rb_DC**) formed a rolled morphology with lamellar ordering, although it lacked the characteristic pH-responsive extension (Fig. 2*A* and *B*, Fig. 3*A* and *E*, and *SI Appendix*, Fig. S4*F*). Furthermore, none of the knockout mutants exhibited reversible extension, demonstrating that the cooperative presence of all four Reb proteins is required for the extension/contraction (*SI Appendix*, Fig. S4). These results support a proposed assembly-pathway model in which RebD contributes to the formation of an α-helix-rich lamellar-ordering intermediate, RebC promotes this lamellar-ordering process without detectable incorporation into the final architecture, and RebA and RebB provide the structural framework that enables this intermediate to mature into the ordered, actuatable lamellar architecture of Rb_WT (Fig. 5).

**Figure 5.**
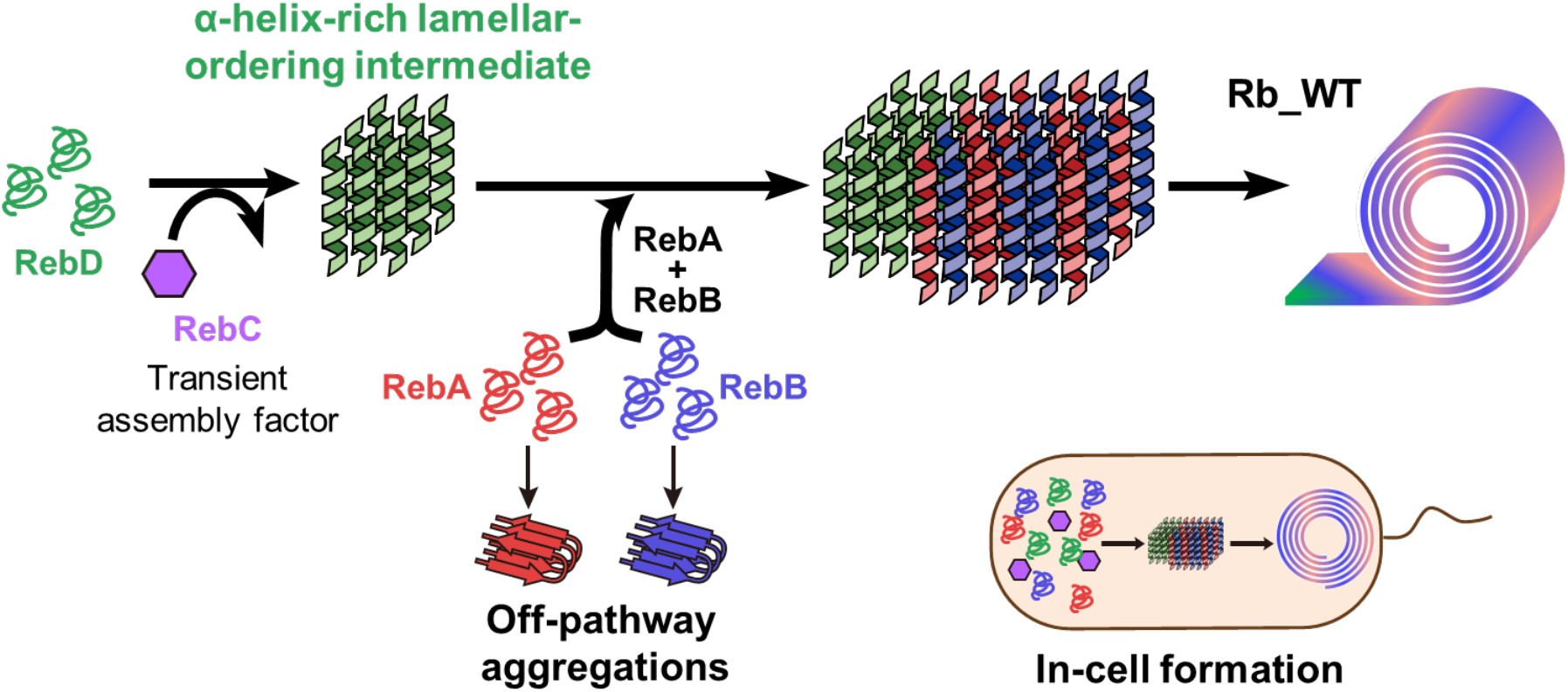
Proposed model for intracellular productive assembly pathway and off-pathway aggregation governed by Reb proteins. R-body formation occurs inside the cell. RebD supports the formation of an α-helix-rich lamellar-ordering intermediate, while RebC promotes this lamellar-ordering process without detectable incorporation into the final R-body. RebA and RebB are subsequently incorporated as major structural components, enabling this intermediate to mature into the ordered, actuatable lamellar architecture of Rb_WT. In the absence of this pathway regulation, Reb proteins are redirected toward β-sheet-rich off-pathway aggregation.

Although RebD was identified in early studies as part of the *reb* locus, its molecular role has remained largely uncharacterized because of its low detectability and unresolved function (23, 27). The absence of RebD (**Rb_ABC**) resulted in globular aggregates with increased β-sheet content and amyloid-like signatures despite the presence of RebA and RebB, indicating that RebA and RebB are prone to aggregation in the absence of RebD (Fig. 3*A*, and *SI Appendix*, Fig. S5*C* and *D*). All mutants that achieved roll formation contained RebD, which was incorporated into Rb_WT as a minor component. Therefore, RebD contributes to productive assembly as a minor incorporated component that supports the formation of an α-helix-rich lamellar-ordering intermediate (Fig. 5). Previous studies have reported that RebA, RebB, and RebD share highly homologous regions (37), suggesting that RebD has structural compatibility with RebA and RebB. This compatibility may induce RebD-containing assemblies to favor α-helical organization of RebA and RebB.

AlphaFold3-predicted models of RebA, RebB, RebC, and RebD suggest that all four Reb proteins have predominantly α-helical structural propensities, consistent with the α-helix-rich character of Rb_WT and **Rb_DC** observed by ATR-FTIR (Fig. 4*A* and *B*, and *SI Appendix*, Fig. S11*A-D*). At the same time, sequence-based aggregation propensity analyses identified aggregation-prone segments in the Reb proteins (*SI Appendix*, Fig. S11*E*), indicating that these proteins may access competing assembly outcomes: productive α-helix-rich lamellar assembly or β-sheet-rich off-pathway aggregation. In this context, the contrasting assembly outcomes of Rb_WT, Rb_DC, and the knockout mutants suggest that RebC and RebD guide the system toward an α-helix-rich lamellar-ordering route, rather than the off-pathway aggregation, thereby establishing a structural precursor to the ordered architecture required for pH-responsive actuation.

Similar pathway-selection behavior has been observed in amphipathic peptides and in the N-terminal segment of huntingtin, although these assemblies eventually convert into β-sheet–rich architectures (38–40). These precedents support the possibility that RebD-containing early assemblies favor an α-helix-rich lamellar-ordering intermediate. Thus, in the absence of RebD, RebA/RebB-containing systems tend toward β-sheet-rich off-pathway aggregation (*SI Appendix*, Fig. S12*D*), whereas RebD supports the formation of α-helical assembly states required for productive R-body formation.

RebC was not detected as a structural component of Rb_WT, yet its absence (**Rb_ABD**) abolished roll formation and promoted aggregation (Fig. 1*E*, Fig. 2*A* and *B*, and *SI Appendix*, Fig. S5*A* and *B*). RebC has previously been proposed to be involved in posttranslational modification of other Reb proteins or translational coupling with RebD, however, these hypotheses were based on electrophoretic analyses or DNA sequences (23, 27). Quantification of roll assemblies per cell showed that mutants containing RebC formed more rolls (Fig. 2*B*). **Rb_ABD** and **Rb_ABC** exhibited similar aggregation behavior, characterized by TEM, ATR-FTIR, and ThT analyses (Fig. 3*A*, and Fig. 4*B* and *C*). Consistently, LC–MS analysis revealed increased levels of aggregation-associated chaperones in these mutants, indicating that the absence of RebC promotes off-pathway aggregation that triggers chaperone production (*SI Appendix*, Fig. S7, and Fig. S12*E*). Therefore, RebC functions as an assembly factor that promotes the lamellar-ordering process without detectable incorporation into the final architecture. Analogous to molecular chaperones that regulate folding versus aggregation (41–43), RebC appears to favor α-helix-rich lamellar ordering over β-sheet-rich off-pathway aggregation (Fig. 5).

RebA and RebB are not required for roll formation but are both essential for pH-responsive extension. Compared to Rb_WT and **Rb_DC**, mutants lacking either RebA (**Rb_BDC**) or RebB (**Rb_ADC**) exhibited decreased roll formation efficiency and increased β-sheet content, indicating that the presence of either RebA or RebB alone shifts the assembly pathway toward β-sheet–rich aggregation, whereas their combined presence or absence suppresses this tendency and enables α-helix-dominant assembly (Fig. 5, and *SI Appendix*, Fig. S12*A-C*). This behavior can be attributed to the intrinsic β-sheet-forming propensity of RebA and RebB as predicted by their sequences (*SI Appendix*, Fig. S11*E*). Western blotting indicates that RebA and RebB form tightly associated hetero-oligomeric complexes in Rb_WT, which were also seen in CBB-stained gels in the previous reports (Fig. 1*E*, ref. 15, 23). These hetero-oligomers, observed only in Rb_WT, are expected to adopt α-helical conformations. Although further structural and biochemical characterization efforts are required, our data suggest that RebA/RebB-containing assemblies acquire α-helix-rich conformations in the presence of RebD and RebC, leading to lamellar organization and functional extension.

In conclusion, this study revealed that all four Reb proteins are required for R-body formation and reversible extension/contraction. Previous studies mainly focused on RebA and RebB as the major structural components of R-bodies, whereas the requirement and molecular functions of RebC and RebD had not been systematically examined. Our results indicate that RebD is incorporated into the wild-type R-body as a minor structural component and RebC is not incorporated into the final architecture in a detectable amount. A series of *reb* gene knockout mutants showed that the presence of both RebC and RebD strongly promotes roll formation. Together, these results support a model in which RebD and the non-incorporated assembly factor RebC establish an α-helix-rich lamellar-ordering intermediate, while RebA and RebB convert this intermediate into the ordered, actuatable architecture of Rb_WT (Fig. 5). Thus, R-body actuation is dictated not simply by the major structural components, but by an assembly pathway that builds nanoscale lamellar order. This study demonstrates that the cooperative interplay of four Reb proteins regulates an assembly pathway that constructs ordered, actuatable R-bodies, providing a conceptual basis for designing dynamic protein architectures whose mechanical responses emerge from pathway-dependent construction of nanoscale order.

## Materials and Methods

### Protein Expression and Isolation

The *E. coli* cells harboring the plasmid of interest were cultured in LB medium until reaching an OD600 of 0.6, followed by the protein expression induced with 1 mM isopropyl β-D-1-thiogalactopyranoside (IPTG) at 37 °C for 18 h. Cell lysis was done by using BugBuster Master Mix. The wild-type strain produced R-body at approximately 600 mg per gram of wet cells. The yields for the **Rb_BDC, Rb_ADC, Rb_ABC**, and **Rb_ABD** mutants were 0.4, 0.3, 0.3, and 0.2 g/g-cell, respectively.

### Liquid Chromatography Mass Spectrometry (LC-MS)

The cell suspensions after expression were boiled at 95 °C for 5 min, followed by freezing and sonication to disrupt the cells. Protein concentrations were adjusted to 25 μg in 50 μL. After reduction and alkylation, the solution was digested with Asp-N, followed by further digestion with trypsin. After digestion, detergents were removed by ethyl acetate/TFA extraction. The aqueous peptide fraction was collected, dried in a centrifugal evaporator, reconstituted in 0.1% TFA/2% acetonitrile, desalted using SDB-XC StageTips, eluted with 0.1% TFA/80% acetonitrile, dried again, and reconstituted in 0.1% TFA/2% acetonitrile for LC–MS/MS analysis. The detailed settings of the LC-MS/MS measurement were the same as reported previously (44). The obtained data were processed on the FragPipe platform using the “LFQ-MBR” workflow with the MSFragger search engine (45) and the IonQuant quantification module (46).

### Ultrathin Sections Preparation for TEM

Cell pellets were pre-fixed with 2.5% glutaraldehyde and 2% paraformaldehyde in 0.1 M PB (pH 7.4), followed by washing three times. Post-fixation was carried out with 1% osmium tetroxide in 0.1 M PB (pH 7.4), followed by a single wash. The samples were then embedded in agarose, dehydrated through a graded dehydration series, and embedded in Epon 812 resin. Polymerization was performed at 60 °C for 48 h. Ultrathin sections with a thickness of 70–80 nm were prepared using an ultramicrotome. The sections were stained with uranyl acetate for 30 min and lead citrate for 15 min prior to observation by TEM.

### TEM Observation and Image Analysis

TEM images were captured using a JEOL1400-Plus electron microscope at 80 kV. The resulting transmission electron microscopy images were analyzed using Fiji (ImageJ, version 1.54p) (47).

### Small-angle X-ray Scattering (SAXS) Measurement

SAXS measurements were performed on a laboratory SAXS system using MicroMax-007HF (Rigaku), a microfocus rotating anode X-ray generator equipped with a Cu-Kα source providing the wavelength λ = 1.54 Å.

### Attenuated Total Reflection Fourier Transform Infrared (ATR-FTIR) Spectroscopy

The measurements were performed using an FT-IR4200 spectrometer (JASCO). Each spectrum was recorded at a resolution of 4 cm^−1^ with the integration of 32 scans.

### Thioflavin T (ThT) fluorescence assays

Thioflavin T (ThT) fluorescence assays were performed using a Cytation 5 plate reader (Agilent BioTek). Fluorescence emission spectra were recorded with an excitation wavelength of 440 nm.

A detailed description of Materials and Methods is available in *SI Appendix*.

## Acknowledgments

We thank Mitsunori Ikeguchi, Tadaomi Furuta, and Hiroka Sugai for helpful discussions. We thank the Integrative Bioscience Facility, Bioscience Center, Research Infrastructure Management Organization, Institute of Science Tokyo, and Ochanomizu Research Facility (ORF), Institute of Science Tokyo for TEM; the Integrative Bioscience Facility at the Institute of Science Tokyo for DNA sequencing; and the Materials Analysis Division, Core Facility Center, Institute of Science Tokyo, for SAXS. We thank the Cell Biology Center Research Core Facility at the Institute of Science Tokyo, for the Q-Exactive mass spectrometry measurements. This work was supported by Japan Society for the Promotion of Science (JSPS) KAKENHI Grant numbers JP25H02254 (T. U.), JP26K17957 (K. K.), JP24KJ1093 (K. D.); the Japan Science and Technology Agency (JST) Adopting Sustainable Partnerships for Innovative Research Ecosystem (ASPIRE) Grant number JPMJAP24B4 (T. U.), ACT-X Grant number JPMJAX25D5 (K. K.).

