## Supporting Information for "Assembly-pathway regulation dictates pH-responsive actuation in the R-body protein machinery"

**This PDF file includes:**

Supporting Text
Supplementary Figures S1 to S12
Supplementary Table S1
Legends for Supplementary Movies S1 and S2
References

**Other supporting materials for this manuscript include the following:**

Movies S1 and S2

**Supporting Text**

**Further Details of Materials and Methods**

**Materials.**
All reagents were purchased from commercial suppliers like Nacalai Tesque and Wako and were used without further purification. The amino acid sequences of proteins used in this study are shown in Table S1. The gene of Rb_WT in a pEX-A2J2 plasmid was purchased from Eurofins. The gene was cloned into pET29b by XbaI and XhoI restriction digests, followed by ligation using Ligation High (Toyobo). The knockout mutants were prepared by inverse PCR using either the KOD Plus Mutagenesis Kit (TOYOBO) or stand-alone reagents consisting of KOD-Plus-Neo (TOYOBO) or PfuTurbo DNA Polymerase, together with FG-DpnI (FastGene) and Ligation High Ver.2 (TOYOBO). The plasmid was transformed into *E. coli* DH5α cells (Toyobo) and purified by QIAprep Spin Miniprep kit (Qiagen). The DNA sequence of the resulting plasmid was verified by DNA sequencing.

**Protein Expression and Isolation.**
Plasmids containing Rb_WT or mutants were transformed into *E. coli* C43 cells. The cells were cultured in LB medium until reaching an OD600 of 0.6, and protein expression was induced with 1 mM isopropyl β-D-1-thiogalactopyranoside (IPTG) at 37 °C for 18 h. Cell pellets were flash-frozen in liquid nitrogen and subsequently thawed. All subsequent procedures were performed at 4 °C unless otherwise stated. Cells were washed by resuspending in 25 mM Tris-HCl (pH 7.5), 100 mM NaCl, and 2 mM EDTA at a density of approximately 10 w/v%, followed by centrifugation (16,000 × *g*, 10 min) and the removal of supernatant. The pellet was resuspended in BugBuster Master Mix (Millipore) at 10 w/v% and stirred at 800 rpm for 15 min at 25 °C, followed by centrifugation (16,000 × *g*, 20 min). The supernatant was discarded, and this step was repeated twice. The pellet was then resuspended in a 10-fold diluted BugBuster Master Mix, centrifuged (5,000 × *g*, 15 min), and the supernatant was removed. This step was repeated three times. The pellet was further resuspended in a 10-fold diluted BugBuster Master Mix (10 mL per gram of cells), followed by centrifugation (16,000 × *g*, 20 min) at 4 °C and the removal of supernatant. Finally, the pellet was resuspended in 10 mL of distilled water and stored at 4 °C. The wild-type strain produced R-body at approximately 600 mg per gram of wet cells. The yields for the **Rb_BDC**, **Rb_ADC**, **Rb_ABC**, and **Rb_ABD** mutants were 0.4, 0.3, 0.3, and 0.2 g/g-cell, respectively.

**SDS-PAGE and Western Blotting.**
Samples were diluted in SDS sample buffer (50 mM Tris-HCl at pH 6.8, 1% SDS, 20% glycerol, 0.01% bromophenol blue, 100 mM DTT) and then heated at 95 °C for 10 minutes. The proteins were separated by SDS-PAGE and subsequently transferred onto polyvinylidene difluoride membranes. The membranes were blocked using 2.5% skim milk in Tris-buffered saline with 0.2% Tween-20. Mouse monoclonal anti-His-tag antibody (clone 9C11, Wako) was used as the primary antibody at a 1:5,000 dilution. Secondary antibodies, HRP-conjugated anti-mouse IgG (Proteintech) were used. Blotted membranes were detected with a chemiluminescence imager (Amersham Imager 680, Cytiva). Coomassie dye staining was applied to the gel using CBB staining solution (Nacalai).

**Phase-Contrast Microscopy Observation of R-body Extension and Contraction in a Flow Channel.**
Phase-contrast images were acquired using an inverted microscope (ECLIPSE Ts2R, Nikon) equipped with a CFI Plan Fluor DLL 100× oil objective (NA 1.30, Ph3, oil immersion; Nikon) and a 4K-UVC digital camera (Relion). For experiments tracking Rb_WT or mutants extension and contraction by phase-contrast microscopy, a 0.5-mm-wide flow channel was created by attaching double-sided tape to a glass slide. Isolated Rb_WT or mutants (OD600 = 0.6) suspended in PBS were introduced into the channel by placing a 10 µL drop at one end, which filled the channel by capillary action. To exchange the solution with a pH 7.0 solution (10 mM Tris-HCl, pH 7.0, 100 mM KCl), a piece of paper was placed at one end of the channel, and 30 µL of solution was then applied to the opposite end so that the solution was wicked through the channel. To test the extension function of Rb_WT and mutants, the solution was replaced with either a pH 3.0 solution (10 mM

acetic acid, pH 3.0, 100 mM KCl) or a pH 5.0 solution (10 mM MES, pH 5.0, 100 mM KCl) using the same procedure. To test the contraction function, the solution was subsequently replaced with a pH 7.0 solution using the same procedure.

**Liquid Chromatography Mass Spectrometry (LC-MS).**
Rb_WT- or mutants-expressing *E. coli* cell pellets were prepared as described in the method section of **Protein Expression and Isolation**. The collected cell pellets were resuspended in PTS buffer (12 mM sodium deoxycholate, 12 mM sodium N-lauroyl sarcosinate, 0.1 M Tris at pH 9.0) and boiled at 95 °C for 5 min. The solution was then frozen at −80 °C for 10 min, followed by sonication at room temperature for 20 min to further disrupt the cells. After cell disruption, protein concentrations were determined using a BCA Protein Assay Kit (Thermo Fisher Scientific, Waltham, MA, USA) and adjusted to 25 µg in 50 µL with PTS buffer. The total protein was reduced with 10 mM dithiothreitol at room temperature for 30 min and alkylated with 50 mM iodoacetamide in the dark at room temperature for 30 min. Following reduction and alkylation, the solution was diluted fivefold with 50 mM ammonium bicarbonate and digested with Asp-N at a ratio of 1:50 (enzyme-to-protein, w/w) at room temperature for 3 h. The peptide fragments were further digested with trypsin at a 1:50 ratio (enzyme-to-protein, w/w) at 37 °C overnight. After digestion, an equal volume of ethyl acetate and 1/20 volume of 10% trifluoroacetic acid (TFA) were added to the peptide solution, followed by vigorous mixing. The mixture was centrifuged at 15,700 × g for 2 min, and the upper organic phase containing detergents was discarded. The lower aqueous phase was dried in a centrifugal evaporator. The peptides were reconstituted in 0.1% TFA and 2% acetonitrile, desalted using StageTips packed with SDB-XC Empore disks (3M, Maplewood, MN, USA), and eluted with 0.1% TFA and 80% acetonitrile. The eluates were dried again and reconstituted in 0.1% TFA and 2% acetonitrile for LC–MS/MS analysis. Peptide samples were analyzed using a Q-Exactive tandem mass spectrometer and an Easy-nLC1000 nanoflow liquid chromatography system (Thermo Fisher Scientific) in the data-dependent acquisition (DDA) mode. The detailed settings of the LC-MS/MS measurement were the same as reported previously (1). The obtained data were processed on the FragPipe platform using the “LFQ-MBR” workflow with the MSFragger search engine (2) and the IonQuant quantification module (3). The amino acid sequence database of *E. coli* proteins for the peptide search was obtained from the UniProt database (https://www.uniprot.org/, downloaded on September 28, 2023). Downstream data processing and statistical analysis were performed using in-house scripts in R.app (ver. 4.5.2).

**Negative-Staining Sample Preparation for Transmission Electron Microscopy (TEM).**
5 µL of the sample was loaded for 1 minute onto a carbon-coated copper grid (U1013, EM Japan), then the excess was removed, and the grid was washed twice with pH 7.0 solution (50 mM citrate–phosphate, pH 7.0), or pH 3.0 solution (10 mM acetic acid, pH 3.0, 100 mM KCl). The sample was negatively stained twice with 5 µL of 1% methylamine tungstate (Nanoprobes, CAS No. 55979-60-7) for 1 min each. After staining, the sample was dried with absorbent paper.

**Ultrathin Sections Preparation for TEM.**
Cell pellets were resuspended in 0.1 M phosphate buffer (PB; pH 7.4) and incubated at 4 °C. The samples were then pre-fixed with 2.5% glutaraldehyde and 2% paraformaldehyde in 0.1 M PB (pH 7.4) at room temperature for 2 h. After pre-fixation, the samples were washed three times with 0.1 M PB (pH 7.4) at 4 °C. Post-fixation was carried out with 1% osmium tetroxide in 0.1 M PB (pH 7.4) at 4 °C for 2 h, followed by a single wash with Milli-Q water at 4 °C. The samples were then embedded in agarose, dehydrated through a graded dehydration series, and embedded in Epon 812 resin. Polymerization was performed at 60 °C for 48 h. Ultrathin sections with a thickness of 70–80 nm were prepared using an ultramicrotome. The sections were stained with uranyl acetate for 30 min and lead citrate for 15 min prior to observation by TEM.

**TEM Observation and Image Analysis.**
TEM images were captured using a JEOL1400-Plus electron microscope at 80 kV. The resulting transmission electron microscopy images were analyzed using Fiji (ImageJ, version 1.54p) (4). Radial averaging of regions of roll architecture was performed using the Radial Profile Extended plugin in Fiji/ImageJ.

**Small-angle X-ray Scattering (SAXS) Measurement.**
SAXS measurements were performed on a laboratory SAXS system using MicroMax-007HF (Rigaku), a microfocus rotating anode X-ray generator equipped with a Cu-Kα source providing the wavelength $\lambda$ = 1.54 Å. The scattered X-ray intensity was collected by the detector Pilatus 100K-S (Rigaku). The sample-to-detector distance was set to 706.5 mm. The samples were resuspended in a relatively high concentration and filled into a 1.5 mm-thick cell. 1D scattering curves *I(q)*, a function of the modulus of the scattering vector $q$, were mainly used for analysis on the Smartlab Studio II software (Rigaku).

**SAXS Analysis.**
Lamellar diffraction peaks in the SAXS profiles were fitted with pseudo-Voigt functions to determine their peak positions and full widths at half maximum (FWHMs). Lamellar spacing was obtained from the fitted peak positions using the relation $2\pi/q_n = d/n$, where $q_n$ is the peak position of the $n$-th order reflection. Specifically, $2\pi/q_n$ was plotted against the peak order $1/n$, and the lamellar spacing $d$ was calculated from the slope of the linear regression. The FWHMs obtained from the pseudo-Voigt fitting were used to evaluate peak broadening associated with structural disorder in the lamellar stacking. To analyze the scattering peaks of spiral state Rb_WT, we used a hollow-cylinder model implemented in SasView software version 6.0.0 (5). The spiral was simulated as a hollow cylinder with 11 nm of wall thickness, 10 µm of length, and 70 nm of radius.

**Attenuated Total Reflection Fourier Transform Infrared (ATR-FTIR) Spectroscopy.**
The measurements were performed using an FT-IR4200 spectrometer (JASCO). Each spectrum was recorded at a resolution of 4 $cm^{-1}$ with the integration of 32 scans. Samples were prepared by suspending isolated Rb_WT and knockout mutants in 50 mM Tris-HCl buffer, pH 7.5, or 50 mM sodium acetate buffer, pH 5.0, followed by centrifugation to remove the supernatant, thereby yielding a concentrated suspension. The suspensions were deposited onto the ATR crystal and air-dried at room temperature before measurement. Before sample measurement, a background spectrum was collected in air and subtracted from each sample spectrum.

**Thioflavin T (ThT) Fluorescence Assays.**
Thioflavin T (ThT) fluorescence assays were performed using a Cytation 5 plate reader (Agilent BioTek). Rb_WT and knockout mutants were suspended in 50 mM Tris-HCl buffer at pH 7.5, and the sample concentration was adjusted to an OD600 of 0.2. Each sample suspension was mixed with 200 µM ThT solution at a 9:1 ratio before measurement. Fluorescence emission spectra were recorded with an excitation wavelength of 440 nm. The fluorescence spectrum was corrected by subtracting the background fluorescence of ThT measured in the absence of protein under identical buffer conditions.

**Prediction of the Aggregation-prone Regions.**
To evaluate the aggregation tendency of each Reb protein, we used four prediction programs: Aggrescan (6), PASTA 2.0 (7), TANGO (8), and WALTZ (9). Each program estimates residue-specific aggregation propensity based on its own calculation method. The parameters for each program were set as follows. For TANGO, the temperature was set to 309.15 K, the ionic strength to 0.02 M, the protein concentration to 1 M, and the pH to 7.0. For WALTZ, the analysis was performed in the high-sensitivity mode at pH 7.0. Aggrescan was used with default settings. For PASTA 2.0, the specificity was set to 90%, and the top pairing energy was 22. Residues with high aggregation propensity were defined as follows. Residues were defined as aggregation-prone when they exceeded the threshold values specified by each program: aggregation-propensity values above −0.02 in Aggrescan, PASTA energy units below −2.8 in PASTA, β-sheet aggregation values above 0.0 in TANGO, and total sequence scores above 0.0 in WALTZ. In this study, an aggregation score of 0 indicates that none of the four programs predicted the residue to be aggregation-prone, whereas a score of 4 indicates that all programs identified the residue as aggregation-prone.

**Reb protein sequence analyses.**

Pairwise alignment of RebA, RebB, and RebD was performed using the Needleman–Wunsch algorithm (10) implemented in EMBOSS Needle (11) available at the EMBL-EBI Job Dispatcher (12). Alignments were performed using the following parameters: substitution matrix = BLOSUM62 (13); gap open = 10; gap extend = 0.5; end gap open = 10; end gap extend = 0.5.

**Supplementary Figures**

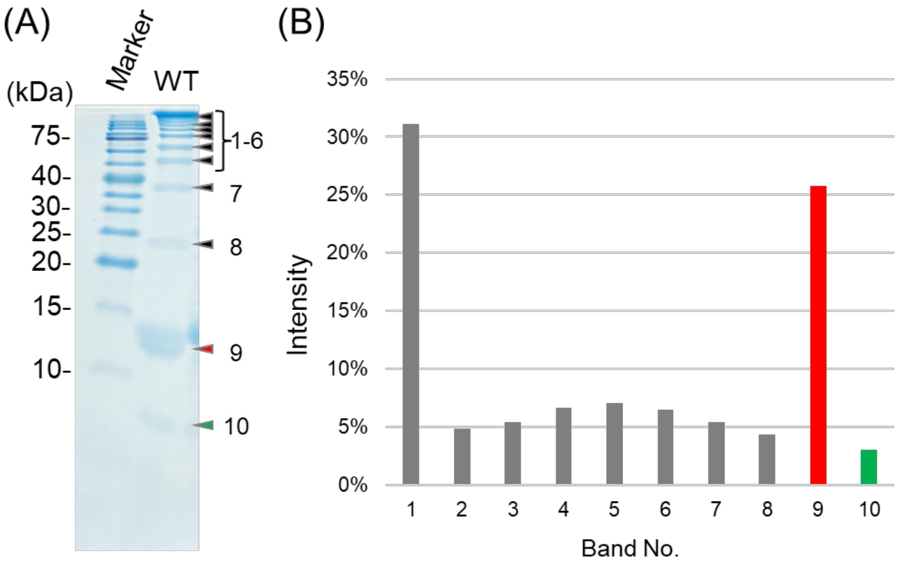


**Fig. S1.** Assignment and relative abundance of the SDS–PAGE bands of isolated Rb_WT. (A) Representative CBB-stained 20% SDS–PAGE gel of isolated Rb_WT. The major bands are numbered from top to bottom. (B) Relative intensity of each band. Bands 1–8 (gray) contain RebA and RebB, band 9 (red) corresponds to RebA, and band 10 (green) corresponds to RebD.

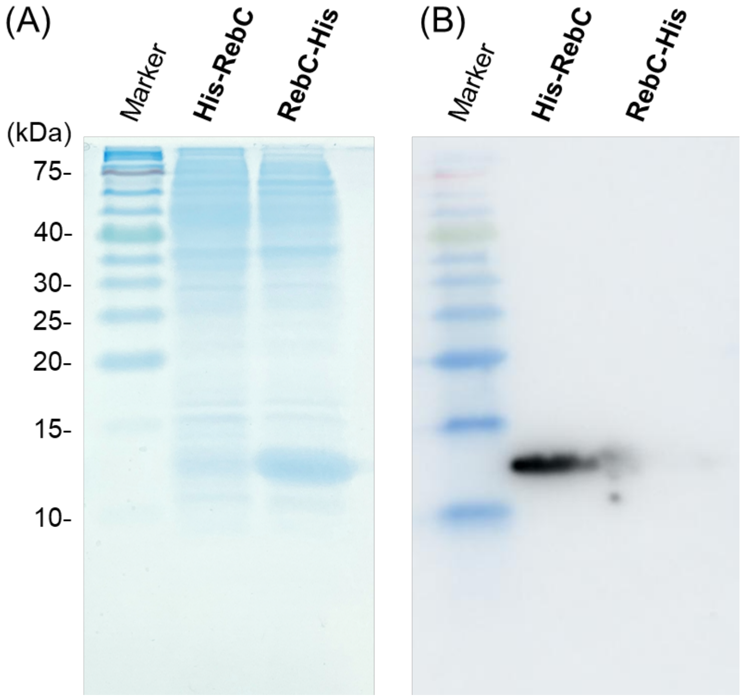


**Fig. S2.** Western blotting analysis of singly expressed RebC fused to an oligo-histidine tag at either the N- or C-terminal. To investigate the reason for no detection of **RebC-His** by Western blotting when using **Rb_WT(His-RebC)** and **Rb_WT(RebC-His)**, RebC with an N-terminal His tag (**His-RebC**) or a C-terminal His tag (**RebC-His**) was expressed individually. (A) SDS–PAGE analysis of cell lysates confirmed the expression of both **His-RebC** and **RebC-His**. (B) Western blotting detected **His-RebC** but not **RebC-His**.

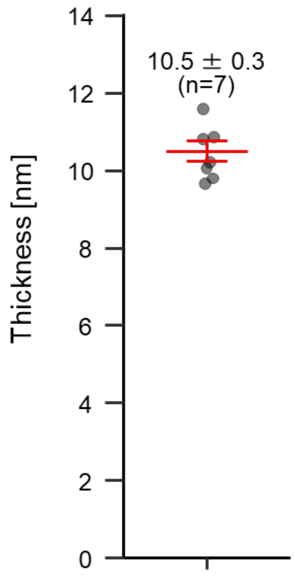


**Fig. S3.** Sheet thickness of Rb_WT measured from individual rolls in TEM images of chemically fixed ultrathin sections. Bars represent mean values with standard error of the mean (s.e.), and individual data points are shown as circles.

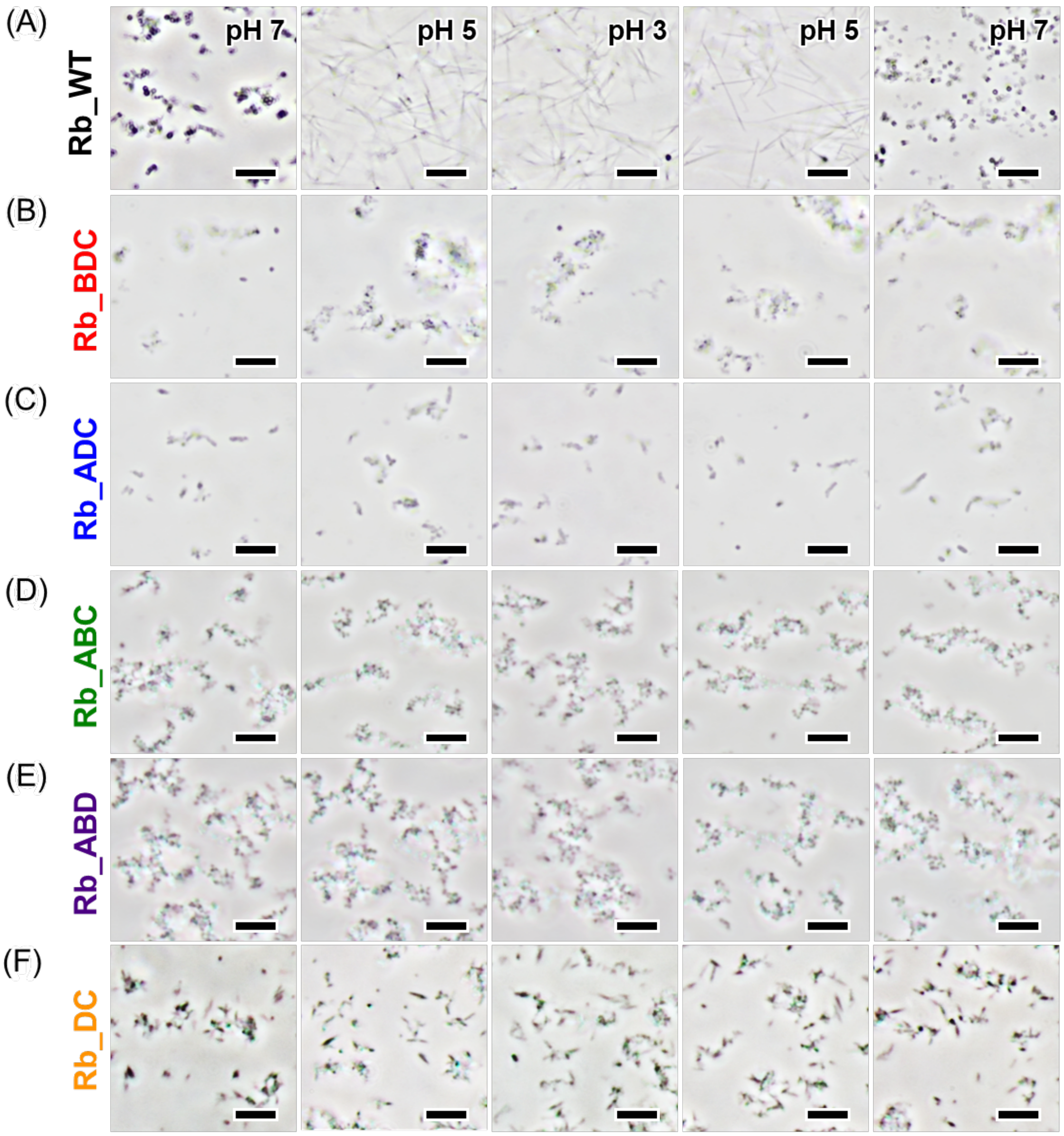


**Fig. S4.** pH-dependent dynamics of Rb_WT and knockout mutants observed by phase-contrast microscopy. (A) Rb_WT showed reversible extension and contraction in response to solution exchange from pH 7.0 to pH 5.0, further to pH 3.0, and back to pH 7.0. (B–F) **Rb_BDC** (B), **Rb_ADC** (C), **Rb_ABC** (D), **Rb_ABD** (E), and **Rb_DC** (F) mutants did not exhibit extension at either pH 5.0 or pH 3.0, nor re-contraction upon return to pH 7.0. Scale bars: 5 μm.

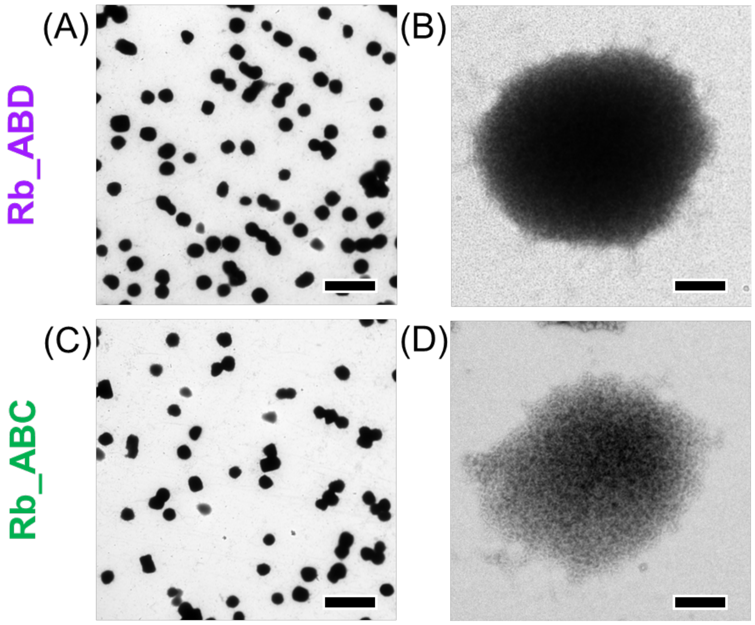


**Fig. S5.** TEM analysis of **Rb_ABD** and **Rb_ABC**. (A-D) TEM images of isolated **Rb_ABD** (A, B), and **Rb_ABC** (C, D) showed globular aggregates. Scale bars: 2 µm (A, C), 200 nm (B, D).

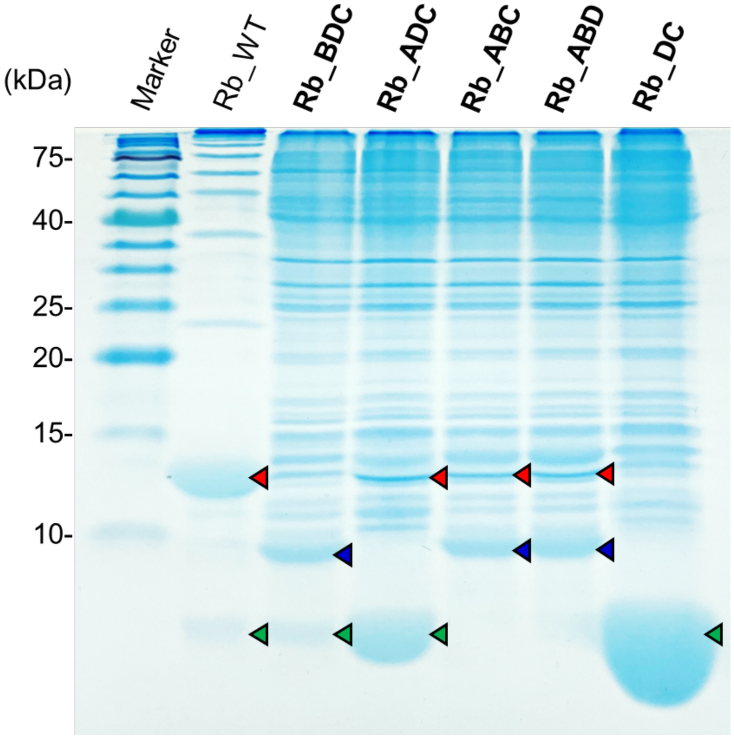


**Fig. S6.** SDS–PAGE analysis of isolated Rb_WT and knockout mutants. After the samples were treated with 1% SDS, 0.1 M DTT, and heated at 95 °C for 10 minutes, Rb_WT showed ladder-like bands on 20% SDS-PAGE gels. In the knockout mutants, bands corresponding to each Reb protein are marked with colored arrowheads: RebA (red, 11.6 kDa), RebB (blue, 11.0 kDa), and RebD (green, 8.5 kDa). RebC (12.5 kDa) was not detected in all samples.

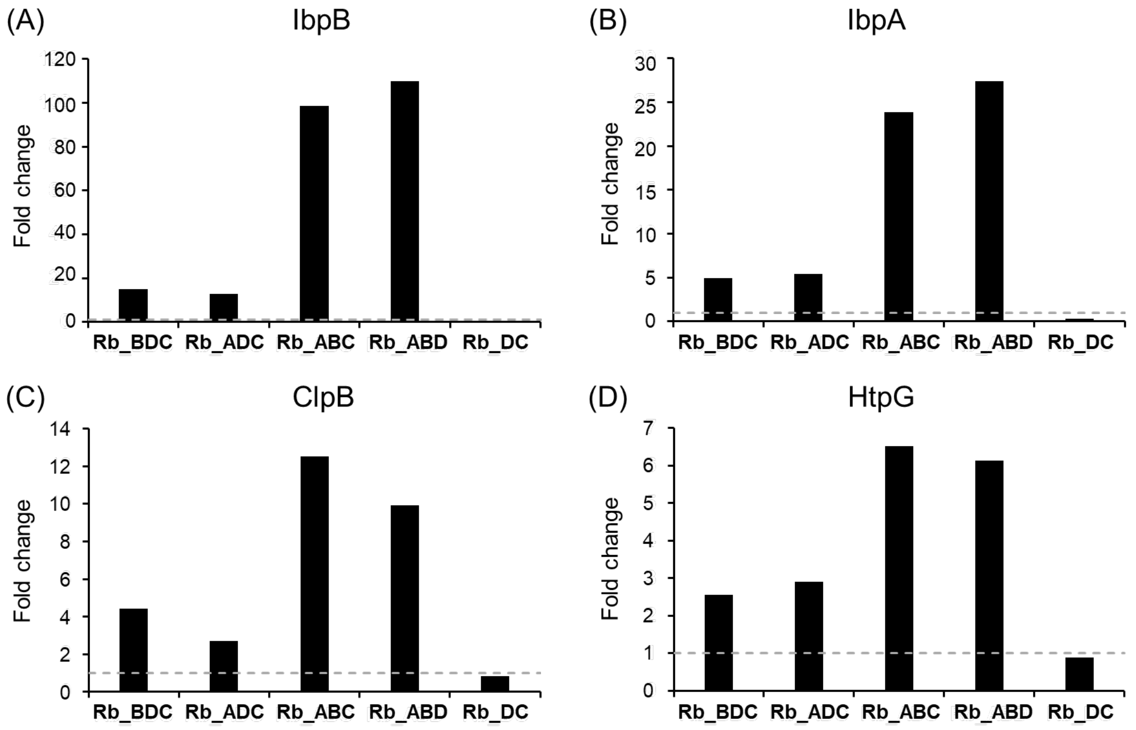


**Fig. S7.** Relative abundance of chaperone proteins in cell lysates of knockout mutants. Fold change in the abundance of chaperone proteins in cell lysates expressing knockout mutants (**Rb_BDC**, **Rb_ADC**, **Rb_ABC**, **Rb_ABD** and **Rb_DC**) relative to the Rb_WT, determined by LC–MS analysis. A fold change of 1 is marked by the horizontal dashed line.

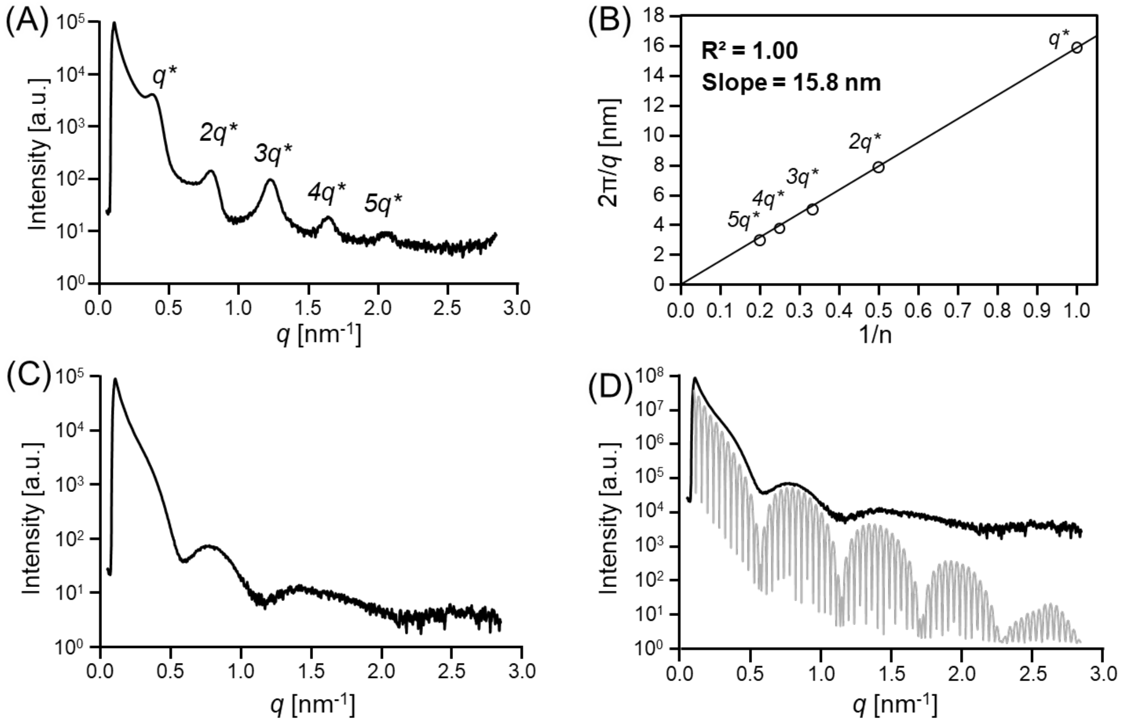


**Fig. S8.** SAXS analysis of spiral and rolled state Rb_WT. (A) SAXS profile of Rb_WT in the rolled state in citrate–phosphate buffer at pH 6.8. (B) Linear relationship between $2\pi/q$ and 1/n for the $2q^*$ to $5q^*$ peaks observed in the SAXS profile of the rolled state Rb_WT. (C) SAXS profile of Rb_WT in the spiral state in citrate–phosphate buffer at pH 3.0. (D) Comparison of the experimental SAXS profile shown in (C) (black line) with the simulated X-ray scattering profile of the hollow cylinder model structure (gray line).

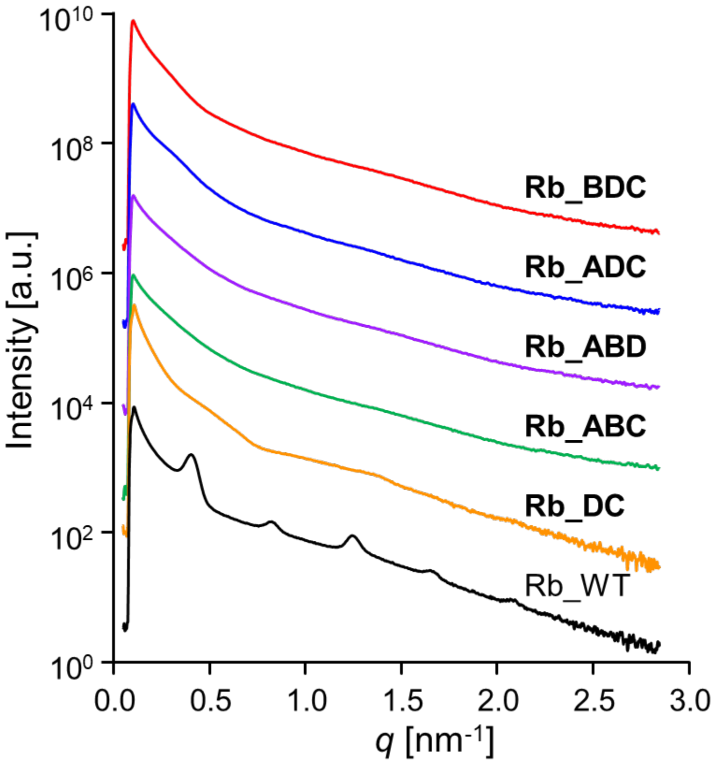


**Fig. S9.** SAXS profiles of *Escherichia coli* expressing Rb_WT or knockout mutants. Cells are suspended in PBS at pH 7.4.

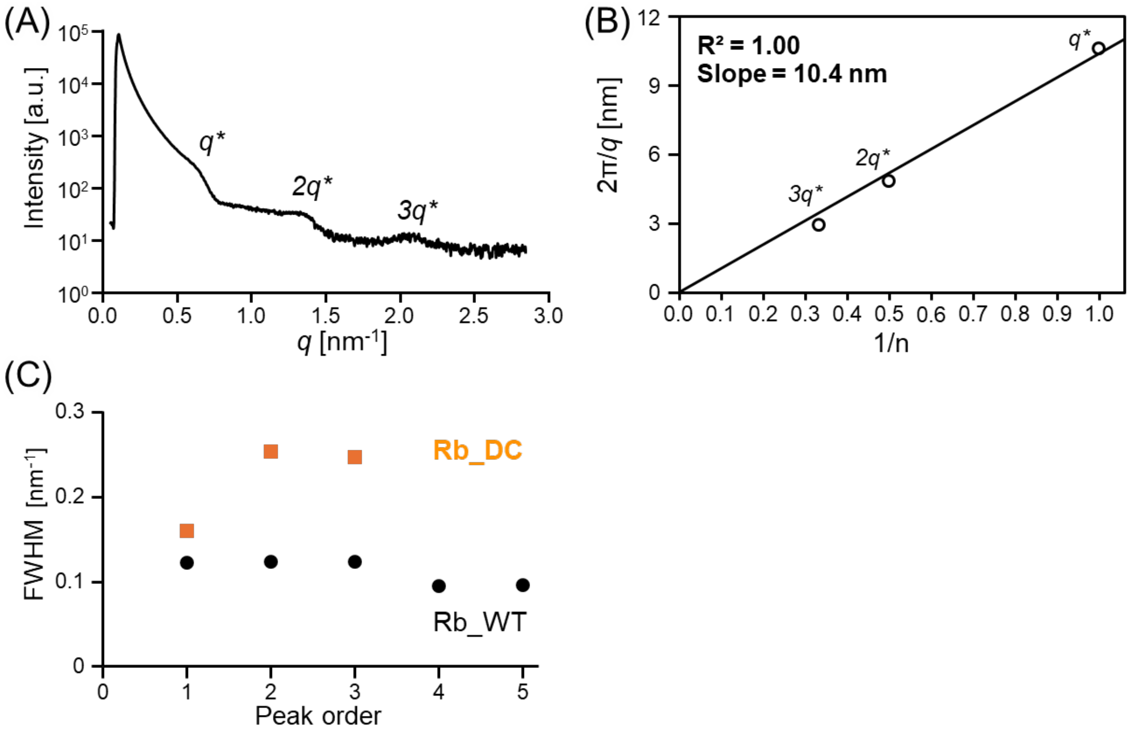


**Fig. S10.** SAXS analysis of **Rb_DC**. (A) SAXS profile of **Rb_DC** in citrate–phosphate buffer at pH 6.8. (B) Linear relationship between $2\pi/q$ and 1/n for the $2q^*$ to $3q^*$ peaks observed in the SAXS profile of the rolled state **Rb_DC**. (C) The full widths at half maximum (FWHMs) of the profiles in (A) as a function of peak order.

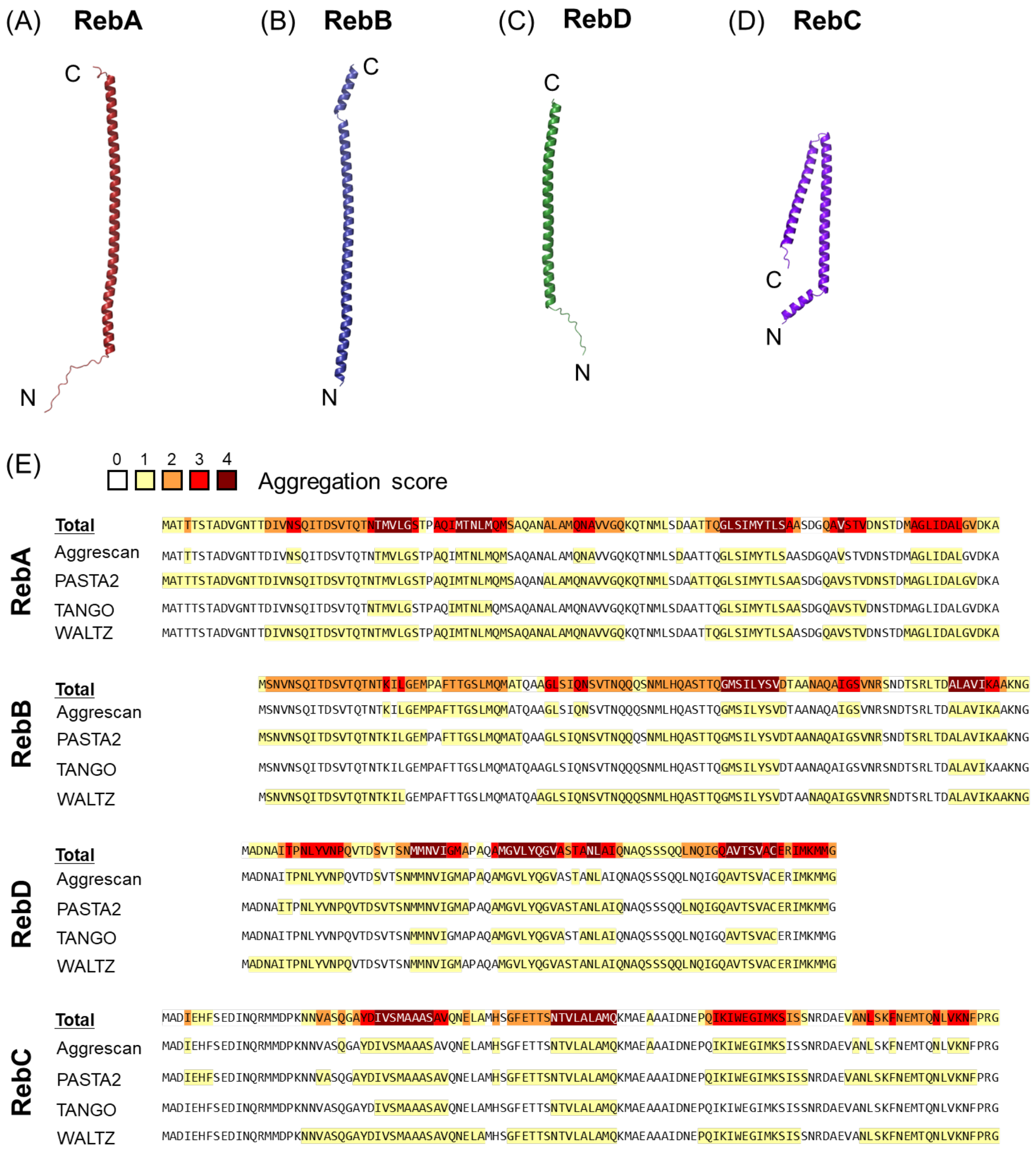


**Fig. S11.** Structure prediction of Reb proteins using AlphaFold3 and aggregation propensity analysis of Reb proteins. (A-D) Monomeric structure predictions of RebA (A), RebB (B), RebD (C), and RebC (D) generated using AlphaFold3 (14). (E) Aggregation propensity analysis of Reb proteins based on four independent prediction programs (Aggrescan (6), PASTA 2.0 (7), TANGO (8), and WALTZ (9)). The sequences are presented with homologous sequences aligned by Clustal Omega (15). Each Reb protein was colored according to its predicted aggregation score (0-4, as indicated in the figure).

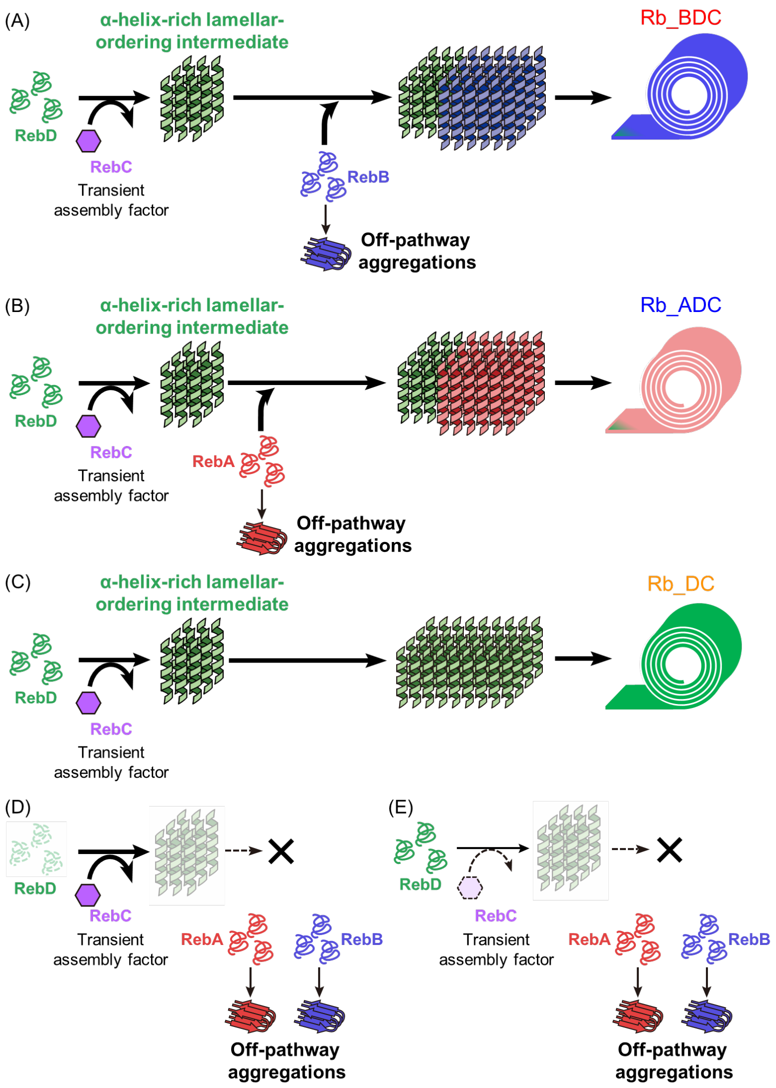


**Fig. S12.** Proposed assembly pathways of knockout mutants. (A, B) In mutants containing RebD, RebC, and one of the two major structural proteins, RebD is proposed to support the formation of an α-helix-rich lamellar-ordering intermediate, while RebC promotes this lamellar-ordering process. This process is followed by incorporation of either RebB (A, **Rb_BDC**) or RebA (B, **Rb_ADC**), leading to rolled architectures that lack the ordered lamellar architecture and pH-responsive extension observed in Rb_WT. (C) In **Rb_DC**, RebD and RebC alone are sufficient to support roll formation; in this mutant, RebD is proposed to serve as the principal structural component of the rolled architecture, whereas RebC likely acts as an assembly factor rather than a stable structural component. (D, E) In contrast, when either RebD or RebC is absent, productive roll assembly does not proceed. Instead, RebA and RebB undergo off-pathway aggregation rather than being incorporated into ordered rolled structures. These models suggest that both RebD and RebC are required to suppress aggregation of RebA/RebB and to direct productive roll assembly.

### Supplementary Tables

**Table S1.** The amino acid sequences of all Reb proteins used in this study

| Protein[a] | Length | MW[b] | Sequence |
|---|---|---|---|
| RebA | 114 | 11622.97 | MATTTSTADVGNTTDIVNSQITDSVTQTNTMVLGSTPA QIMTNLMQMSAQANALAMQNAVVGQKQTNMLSDAAT TQGLSIMYTLSAASDGQAVSTVDNSTDMAGLIDALGV DKA |
| RebB | 105 | 10965.21 | MSNVNSQITDSVTQTNTKILGEMPAFTTGSLMQMATQ AAGLSIQNSVTNQQQSNMLHQASTTQGMSILYSVDTA ANAQAIGSVNRSNDTSRLTDALAVIKAAKNG |
| RebC | 114 | 12523.12 | MADIEHFSEDINQRMMDPKNNVASQGAYDIVSMAAAS AVQNELAMHSGFETTSNTVLALAMQKMAEAAAIDNEP QIKIWEGIMKSISSNRDAEVANLSKFNEMTQNLVKNFP RG |
| RebD | 81 | 8491.75 | MADNAITPNLYVNPQVTDSVTSNMMNVIGMAPAQAM GVLYQGVASTANLAIQNAQSSSQQLNQIGQAVTSVACE RIMKMMG |
| RebA-His | 124 | 12704.05 | MATTTSTADVGNTTDIVNSQITDSVTQTNTMVLGSTPA QIMTNLMQMSAQANALAMQNAVVGQKQTNMLSDAAT TQGLSIMYTLSAASDGQAVSTVDNSTDMAGLIDALGV DKAGGGSHHHHHH |
| RebB-His | 115 | 12046.29 | MSNVNSQITDSVTQTNTKILGEMPAFTTGSLMQMATQ AAGLSIQNSVTNQQQSNMLHQASTTQGMSILYSVDTA ANAQAIGSVNRSNDTSRLTDALAVIKAAKNGGGGSHH HHHH |
| RebC-7His | 125 | 13741.34 | MADIEHFSEDINQRMMDPKNNVASQGAYDIVSMAAAS AVQNELAMHSGFETTSNTVLALAMQKMAEAAAIDNEP QIKIWEGIMKSISSNRDAEVANLSKFNEMTQNLVKNFP RGGGGSHHHHHHH |
| RebC-6His | 124 | 13604.20 | MADIEHFSEDINQRMMDPKNNVASQGAYDIVSMAAAS AVQNELAMHSGFETTSNTVLALAMQKMAEAAAIDNEP QIKIWEGIMKSISSNRDAEVANLSKFNEMTQNLVKNFP RGGGGSHHHHHH |
| His-RebC | 124 | 13604.20 | MHHHHHHGGGSADIEHFSEDINQRMMDPKNNVASQG AYDIVSMAAASAVQNELAMHSGFETTSNTVLALAMQK MAEAAAIDNEPQIKIWEGIMKSISSNRDAEVANLSKFNE MTQNLVKNFPRG |
| RebD-His | 91 | 9572.83 | MADNAITPNLYVNPQVTDSVTSNMMNVIGMAPAQAM GVLYQGVASTANLAIQNAQSSSQQLNQIGQAVTSVACE RIMKMMGGGGSHHHHHH |

[a]The Rb_WT expresses RebA, RebB, RebC, and RebD. Each **Rb_WT(RebX-His)** and **Rb_WT(His-RebX)** expresses **RebX-His** instead of RebX and the other 3 Reb proteins remain identical to those in Rb_WT. **RebC-7His** was expressed in **Rb_WT(RebC-His)** and **RebC-6His** was singly expressed.

[b] The values of MW were calculated based on the primary sequence using the ProtParam tool in ExPASy (16).

**Legends for Supplementary Movies S1 and S2**

**Movie S1 (separate file).** Phase contrast microscopy observation of Rb_WT extension by exchanging solution from a pH 7.0 solution (10 mM Tris-HCl, pH 7.0, 100 mM KCl) to a pH 3.0 solution (10 mM acetic acid, pH 3.0, 100 mM KCl).

**Movie S2 (separate file).** Phase contrast microscopy observation of Rb_WT contraction by exchanging buffer from a pH 3.0 solution (10 mM acetic acid, pH 3.0, 100 mM KCl) to a pH 7.0 solution (10 mM Tris-HCl, pH 7.0, 100 mM KCl).
